# *De Novo* Lipid Labeling for Comprehensive Analysis of Subcellular Distribution and Trafficking in Live Cells

**DOI:** 10.1101/2020.10.16.342683

**Authors:** Jun Zhang, Jia Nie, Haoran Sun, John-Paul Andersen, Yuguang Shi

## Abstract

Lipids exert dynamic biological functions which are determined both by their fatty acyl compositions and precise spatiotemporal distributions inside the cell. There are more than 1000 lipid species in a typical mammalian cell. However, it remains a daunting task to investigate any of these features in live cells for each of the more than 1000 lipid species. Here we resolved this issue by developing a *de novo* lipid labeling method for major lipid species, including glycerolipids, glycerophospholipids, and cholesterol esters by using a single fluorescent probe. The method not only allowed us to probe the precise subcellular distribution and trafficking of individual lipid species in live cells, but also uncovered some unexpected biological functions of previously reported lipid metabolic enzymes that were not possible by conventional biochemical methods. We envision that this method will become an indispensable tool for the functional analysis of individual lipid species and numerous lipid metabolic enzymes and transporters in live cells.

## Introduction

Eukaryotic cells contain thousands of different lipid species which are classified into seven categories according to their fatty acyl structures and pathways of biosynthesis, including glycerol lipids, glycerophospholipids, sterol lipids, sphingolipids, prenol lipids, and saccharolipids^1^. Lipids are highly organized within eukaryotic cells with diversified biological activities, such as membrane structure, energy storage, signal transduction, autophagy, epigenetic modifications, and vesicular trafficking^2^. The precise function of an individual lipid is determined by both its acyl composition and subcellular localization. However, in contrast to DNA and proteins, there is a lack of a universal labeling method for individual lipid species, which makes it very difficult to investigate the spatiotemporal distribution of a given lipid molecule preventing the ability to answer the key question as to when a lipid molecule performs each of its functions within a live cell.

Although various attempts have been made in recent years to investigate subcellular localization and trafficking of lipids by microscopic visualization in live cells^2–4^, it has been met with limited success to date due to a lack of acyl-specific labeling of endogenous lipids. Consequently, most lipid trafficking studies reported to date relied heavily on the use of exogenously labeled lipids or nonspecific fluorescent labeling, such as BODIPY for neutral lipids and Filipin for cholesterol, which renders the results difficult to interpret. BODIPY labels all endogenous neutral lipids without any specificity for acyl composition, whereas fluorophore-conjugated lipids do not mimic the biological functions of the endogenous counterparts, since most of them are subjected to lysosomal degradation after endocytic uptake^5^. In addition, none of the existing lipid labeling methods can be used to monitor the subcellular localization and trafficking of endogenous lipids. In this study, we resolved these issues by developing a *de novo* lipid labeling method which can be used for the analysis of precise subcellular localization and trafficking of individual lipid species in live cells. The method is based on the principle that all lipid species undergo a periodical remodeling process after initial synthesis to achieve appropriate acyl contents or to repair damaged acyl chains from oxidative stress^6^. The remodeling process, also known as the Lands cycle, takes place at the ER-mitochondrial junctions, and is catalyzed by a superfamily of phospholipases and acyltransferases^7–9^. The remodeling process provides a unique opportunity to label individual species by feeding the cultured cells with various lyso-phospholipids and a nitro-benzoxadiazolyl (NBD) labeled acyl-CoA (NBD-CoA) with predesignated acyl composition (NBD is a fluorescent probe). Incorporation of NBD-labeled fatty acyl chains into newly remodeled lipids generates green fluorescence which allows for confocal imaging analysis of the subcellular localization and trafficking of lipid molecules based on their precise acyl composition in live cells (Fig. 1a). We have successfully applied this method to investigate the subcellular localization and trafficking of major lipid species with precise acyl compositions, including cholesterol esters, neutral lipids, and phospholipids in live cells by confocal imaging analysis.

**Fig. 1:**
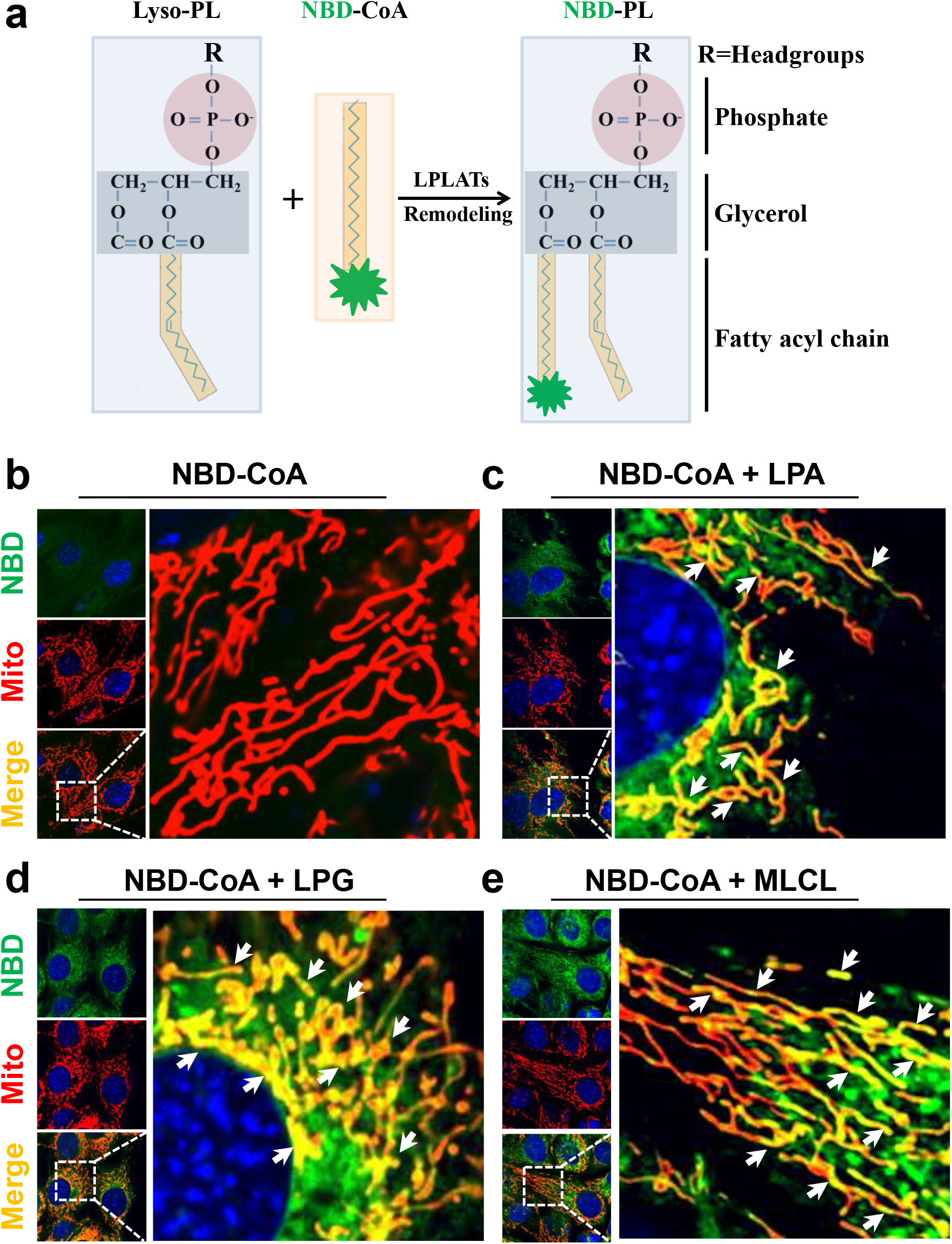
Labeling and confocal imaging of phospholipids in live cells. **a,** Schematic representation of the principle of lipid labeling using NBD-CoA through lipid remodeling. **b-e,** Confocal imaging analysis of newly remodeled and NBD-labeled phospholipids in live C2C12 cells. C2C12 cells were incubated with NBD-palmitoyl-CoA alone (**b**), or NBD-palmitoyl-CoA and LPA (**c**), LPG (**d**), or MLCL (**e**). Mitochondria were stained with MitoTracker-Red. Arrows highlight the co-localization of NBD-PL with mitochondria.

## Results

### An acyl chain-specific *de novo* lipid labeling method to monitor lipid trafficking in live cells

Lipid remodeling is a widespread strategy after the initial synthesis by the Kennedy pathway, which plays a critical role in maintaining appropriate acyl composition of a given lipid species. The remodeling process begins with hydrolysis of fatty acyl chains, followed by reacylation with the required fatty acid, which is catalyzed by a family of acyltransferases using acyl-CoA as the acyl donor (Fig. 1a). We questioned whether the remodeling process could provide an opportunity for *de novo* labeling with selective lipid species by NBD-labeled acyl-CoAs, which can be imaged by confocal microscopy. We first tested this hypothesis for the *de novo* labeling of phosphatidic acids (PA) in C2C12 cells, a skeletal myocyte cell line. PA is a precursor for the synthesis of all the phospholipids, including phosphatidylcholine (PC), phosphatidylethanolamine (PE), phosphatidylserine (PS), phosphatidylinositol (PI), phosphatidylglycerol (PG), and cardiolipin (CL). C2C12 cells were stained with Mitotracker Red, a mitochondria specific fluorescent dye, and cultured in the presence of either NBD-palmitoyl-CoA alone or in the presence of lysophosphatic acid (LPA), a substrate for PA synthesis. No significant green fluorescence signal was detected in cells cultured with NBD-palmitoyl-CoA alone (Fig. 1b). In contrast, supplementation of culturing medium with both NBD-palmitoyl-CoA and LPA resulted in detection of strong fluorescent signals (Fig. 1c), suggesting an acyl specific labeling of newly remodeled PA. PA is primarily remodeled in the endoplasmic reticulum (ER), and is transported to mitochondria where it is used as the precursor for the synthesis of mitochondrial specific phospholipids, including PG and CL. Consistent with this notion, newly labeled PA first appeared in the ER, and part of the NBD-labeled PA was rapidly transported to mitochondria, as evidenced by co-localization of red (mitochondria) and green (PA) (Fig. 1b and 1c, highlighted by arrows), and by results from time-lapse confocal imaging analysis (Extended Data Fig. 1a and 1b). In support of the specificity of the labeling method, incubation of C2C12 cells with NBD-palmitoyl-CoA in the presence of lysophosphatidylglycerol (LPG) or monolysocardiolipin (MLCL), the substrates for the remodeling of PG and CL, also resulted in specific labeling of PG and CL which are rapidly transported to mitochondria (Fig. 1d and 1e, highlighted by arrows). In contrast to CL which is exclusively localized in mitochondria after remodeling (Extended Data Fig. 2, highlighted by arrows), only part of the newly remodeled PC was transported to mitochondria, as shown by the time-lapse confocal imaging analysis (Extended Data Fig. 3, arrows highlight the co-localization of NBD-PC with mitochondria (3a) or ER (3b), respectively).

One of the major concerns for the labeling method is the attachment of NBD to palmitoyl −CoA, which could render the acyl-CoA inactive. Using thin layer chromatography (TLC) analysis, we addressed this concern by validating whether the fluorescent signals we detected from the live C2C12 cells were indeed generated from the remodeled phospholipids. In further support of the specificity of the labeling, the results show that NBD-labeled phospholipids, including PA, PG, and PC can be detected only if both NBD-palmitoyl-CoA and lysophospholipids were added to the culture medium (Extended Data Fig. 4a-c, highlighted by arrow). The results also suggest that attachment of NBD did not affect its biological activity, since the NBD-palmitoyl-CoA was successfully used as the acyl donor by the endogenous acyltransferases.

Using isolated mouse embryonic fibroblasts (MEFs) and primary hepatocytes, we determined whether the lipid labeling method could be applied to study the subcellular localization and traffic of phospholipids in primary cells. Consistent with findings from C2C12 cells, treatment of MEFs and primary hepatocytes with both NBD-palmitoyl-CoA and LPA or LPG resulted in NBD-palmitoyl-CoA dependent remodeling of PA or PG, respectively, which are also promptly translocated to the mitochondria both in MEFs and primary hepatocytes (Extended Data Fig. 5a and 5b), suggesting that the labeling method can also be used to study lipid trafficking in primary cells as well.

### TRIAP1/PRELI are required for mitochondrial transport of PA and PG, but not PC

Currently, analysis of lipid transporter activity is commonly carried out by liposome-based assays *in vitro* using the purified protein to be tested^10^. Though this method provide a tool to investigate the activity of a protein or a protein complex in lipid binding and trafficking *in vitro*, some of the purified proteins, especially the membrane proteins, may not be functional *in vitro*. Also, currently there is no method to directly validate the function of a protein in lipid trafficking in live cells. Additionally, the liposome-based assay does not faithfully mimic the endogenous membrane environment, such as the ER and mitochondrial membranes. We next determined whether the *de novo* lipid labeling technique can be used to identify lipid transporter activity of a given protein in live cells. We first tested the function of the TRIAP1/PRELI protein complex which was previously shown to facilitate the transport of PA from the ER to mitochondria^10^. Consistent with the previously reported function of the complex, ablation of PRELID1 or TRIAP1 in C2C12 cells significantly impaired PA transport from the ER to mitochondria, leading to the ER retention of newly remodeled PA (Fig. 2a-c, highlighted by arrows, Pearson’s correlation coefficient of mitochondria and NBD-PA was quantified in Fig. 2g). The findings are further supported by results from time-lapse confocal imaging analysis in live C2C12 cells stained with Mitotracker red (Extended Data Fig. 6a-c). Surprisingly, relative to the vector controls (Fig. 2d), PRELID1 deficiency, but not TRIAP1, also significantly impaired the transport of remodeled PG to mitochondria (Fig. 2e and 2f, highlighted by arrows, Pearson’s correlation coefficient of mitochondria and NBD-PG was quantified in Fig. 2h and 2i, respectively). This result is consistent with the previous report that PRELI deficiency, but not TRIAP1, reduced the PG in mitochondria^10^. In contrast, neither PRELID1 nor TRIAP1 deficiency had any effect on the transport of remodeled PC to mitochondria in C2C12 cells when cultured in the presence of NBD-palmitoyl-CoA and LPC as the substrates for PC remodeling (Extended Data Fig. 7a-c, Pearson’s correlation coefficient of mitochondria and NBD-PC was quantified in Extended Data Fig. 7d and 7e).

**Fig. 2:**
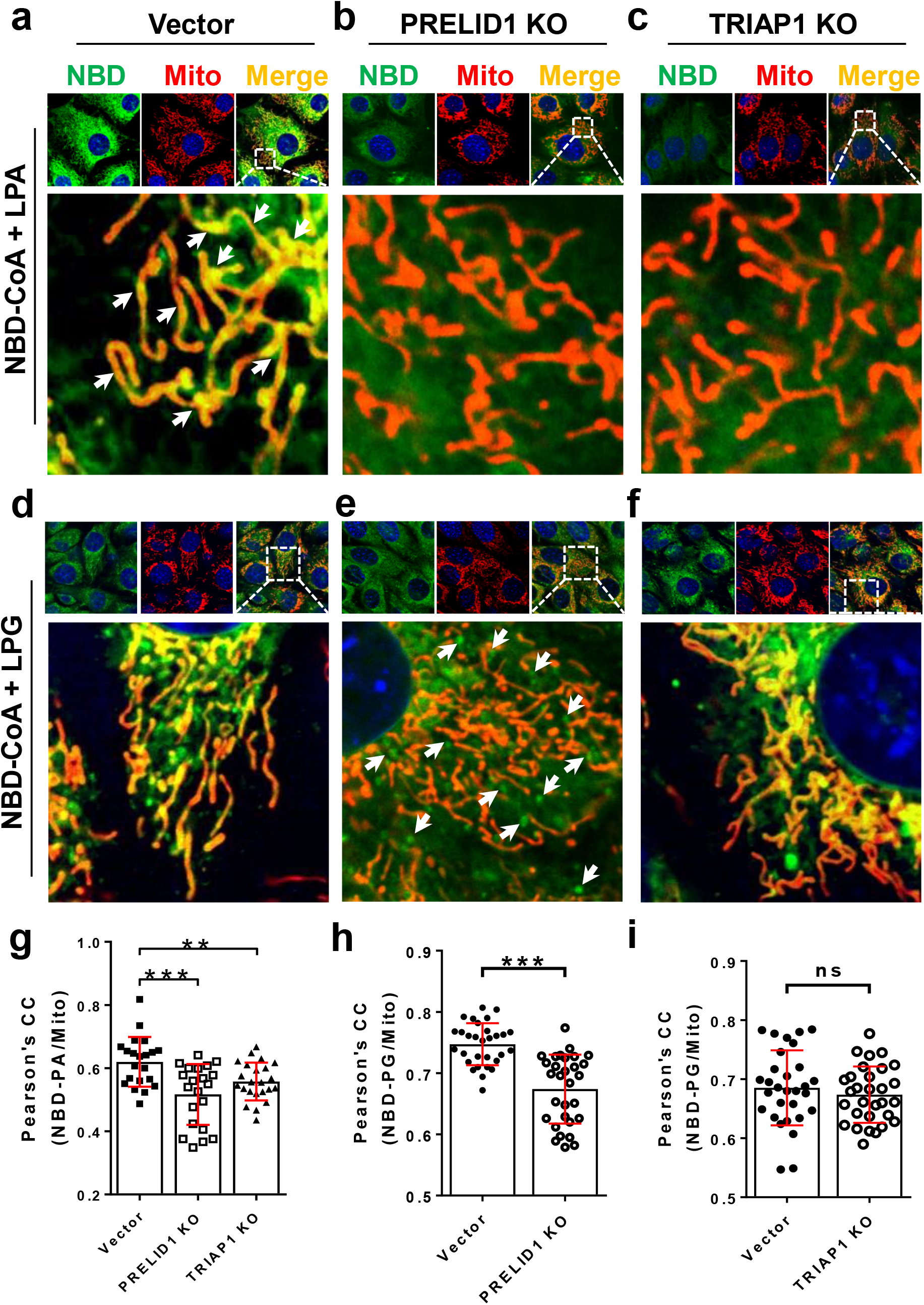
TRIAP/PRELID1 complex is required for mitochondrial PA and PG, but not PC transport. **a-c,** Confocal imaging analysis of the labeling and mitochondrial transport of PA in C2C12-vector control (**a**), PRELID1 KO (**b**) and TRIAP1 KO (**c**) cells. Arrows highlight the co-localization of NBD-PA with mitochondria. **d-e,** Confocal imaging analysis of the labeling and mitochondrial transport of PG in C2C12-vector control (**d**), PRELID1 KO (**e**) and TRIAP1 KO (**f**) cells. Arrows highlight the NBD-PG outside of mitochondria. **g,** Pearson’s correlation coefficient of mitochondria and NBD-PA in vector, PRELID1 KO and TRIAP1 KO cells. n=21-23 cells/group. **h-i,** Pearson’s correlation coefficient of mitochondria and NBD-PG in vector and PRELID1 KO (**h**) or TRIAP1 KO cells (**i**). n=30 cells/group. Cells were starved in KRPH buffer for 1 hr, and then incubated with NBD-palmitoyl-CoA and LPA (**a-c**) or LPG (**d-f**) for 15 min. Mitochondria were stained with MitoTracker-Red. Data are represented as mean ± SD, **p<0.01, ***p<0.001, ns, no significance by student’s t-test.

### An unexpected role of StARD7 in PG transport

The StAR-related lipid transfer domain protein 7 (StARD7) is a member of the START domain-containing family of proteins, and is implicated in transport of PC between membranes^11^. Targeted deletion of StARD7 also causes mitochondrial dysfunction and loss of mitochondrial crista, but the underlying causes remain elusive^12^. Using C2C12 cells with CRISPR/Cas9-mediated deletion of the *StARD7* gene, we investigated the role of StARD7 in mediating phospholipid trafficking between the ER and mitochondria. Consistent with the reported role in PC transport, ablation of StARD7 indeed impaired the transport of newly remodeled PC to mitochondria, as evidenced by a lack of co-localization of red (mitochondria) with green (PC) (Fig. 3b) relative to the vector control (Fig. 3a, Pearson’s correlation coefficient of mitochondria and NBD-PC was quantified in Fig. 3c). Surprisingly, StARD7 is also required for the transport of newly remodeled PG to mitochondria, as evidenced by decreased co-localization with mitochondria (Fig. 3d-e, highlighted by arrows, Pearson’s correlation coefficient of mitochondria and NBD-PG was quantified in Fig. 3f). In contrast, StARD7 is not required for the trafficking and transport of PA from the ER to mitochondria, as StARD7 deficient cells exhibited normal PA trafficking from the ER to mitochondria (Fig. 3g-i).

**Fig. 3:**
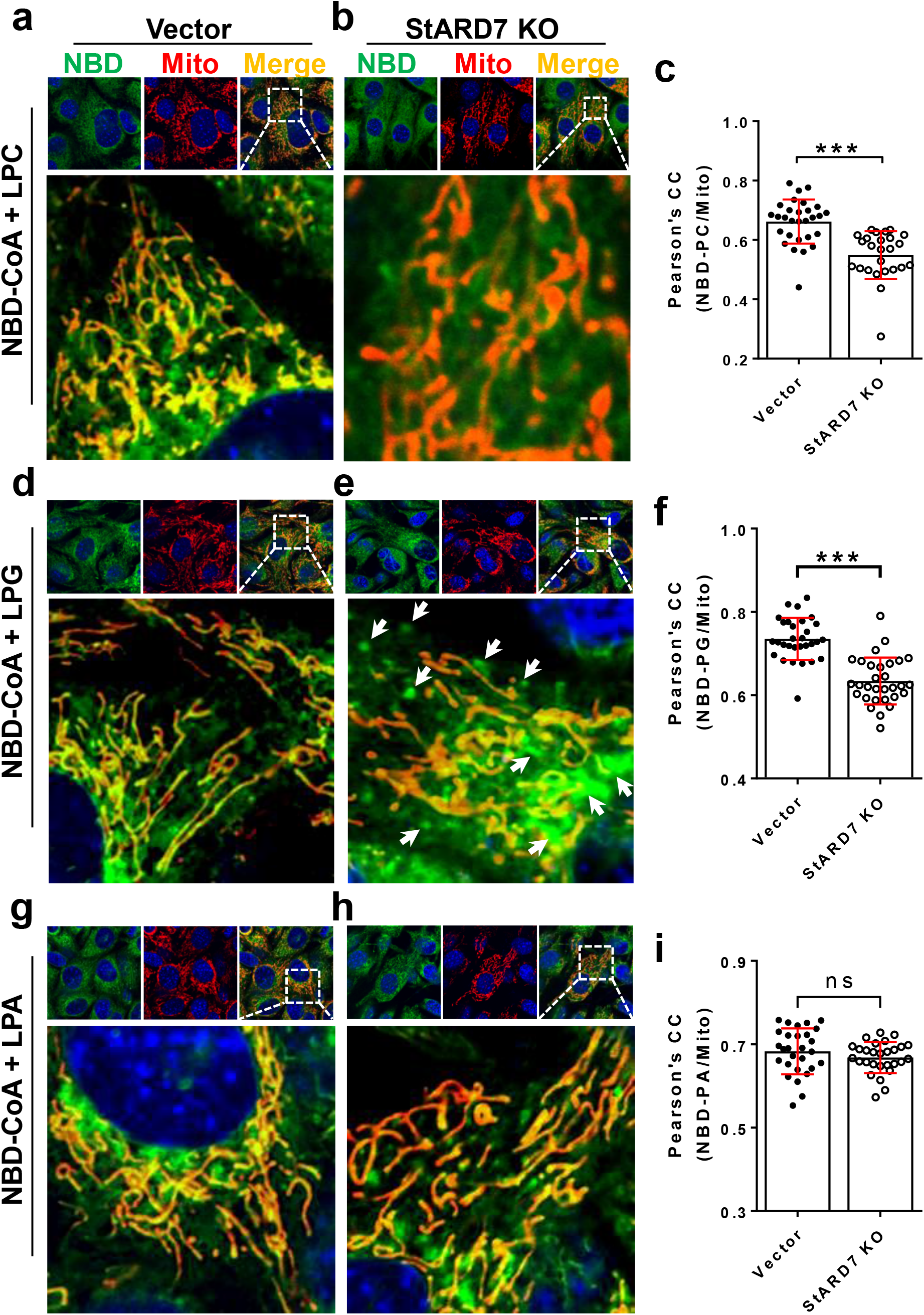
StARD7 is required for mitochondrial PC and PG, but not PA transport. **a-b,** Confocal imaging analysis of the labeling and mitochondrial transport of PC in C2C12-vector control (**a**) and StARD7 KO (**b**) cells. **c,** Pearson’s correlation coefficient of mitochondria and NBD-PC in vector and StARD7 KO cells. N=25-29 cells/group **d-e,** Confocal imaging analysis of the labeling and mitochondrial transport of PG in C2C12-vector control (**d**) and StARD7 KO (**e**) cells. **f,** Pearson’s correlation coefficient of mitochondria and NBD-PG in vector and StARD7 KO cells. N=30 cells/group. **g-h,** Confocal imaging analysis of the labeling and mitochondrial transport of PA in C2C12-vector control (**g**) and StARD7 KO (**h**) cells. **i,** Pearson’s correlation coefficient of mitochondria and NBD-PA in vector and StARD7 KO cells. N=30 cells/group. Cells were starved in KRPH buffer for 1 hr, and then incubated with NBD-palmitoyl-CoA and LPC (**a-b**), LPG (**d-e**), or LPA (**g-h**) for 15 min. Mitochondria were stained with MitoTracker-Red. Data are represented as mean ± SD, ***p<0.001, ns, no significance by student’s t-test.

### A surprising role of ACAT1 in regulating mitochondrial cholesterol transport

Mitochondrial cholesterol transport plays a vital role in the synthesis of steroids, oxysterols, and hepatic bile acids^13, 14^. Mitochondrial membranes are intrinsically low in cholesterol content when compared with other membranes, and therefore rely upon cholesterol transport from other organelles for steroidogenesis. The prevailing hypothesis of mitochondrial cholesterol transport is that hydrolysis of cholesterol esters from lipid droplets provides the major source for mitochondrial cholesterol transport^15^. However, it is not known whether cholesterol esters can also enter the mitochondria. The ER is the major site for cholesterol esterification, as both acylcoenzyme A: cholesterol acyltransferase-1 and 2 (ACAT1 and ACAT2) are localized in the ER^16^. Using the lipid labeling method, we next determined a role of cholesterol acylation by ACATs in the ER on mitochondrial cholesterol transport. We first tested whether NBD-palmitoyl-CoA could be used to label cholesterol esters by culturing C2C12 cells in the presence of NBD-palmitoyl-CoA alone or with cholesterol. Similar to the findings with phospholipids, addition of NBD-palmitoyl-CoA together with cholesterol to the culture medium resulted in the labeling of cholesterol esters with green fluorescence (Fig. 4a and 4b). Surprisingly, part of the cholesterol esters was rapidly transported to mitochondria (Fig. 4b, highlighted by arrows). In support of the specificity of the labeling, treatment of C2C12 cells with avasimibe, a potent inhibitor of both ACAT1 and ACAT2 enzymes, completely abolished the labeling and mitochondrial localization (Fig. 4c, Pearson’s correlation coefficient of mitochondria and NBD was quantified in Fig. 4d). Consistent with the findings, avasimibe also significantly reduced mitochondrial content of cholesterol esters (Fig. 4e), leading to a significant increase in total free cholesterol content in the cytosol (Fig. 4f). In contrast, treatment of etomoxir, an inhibitor of carnitine palmitoyl transferase which is required for mitochondrial fatty acid uptake, did not significantly affect mitochondrial cholesterol transport (Fig. 4g and 4h). The results further confirm that the newly remodeled cholesterol esters, but not NBD-labeled free fatty acids released from the cholesterol ester, were indeed transported to mitochondria.

**Fig. 4:**
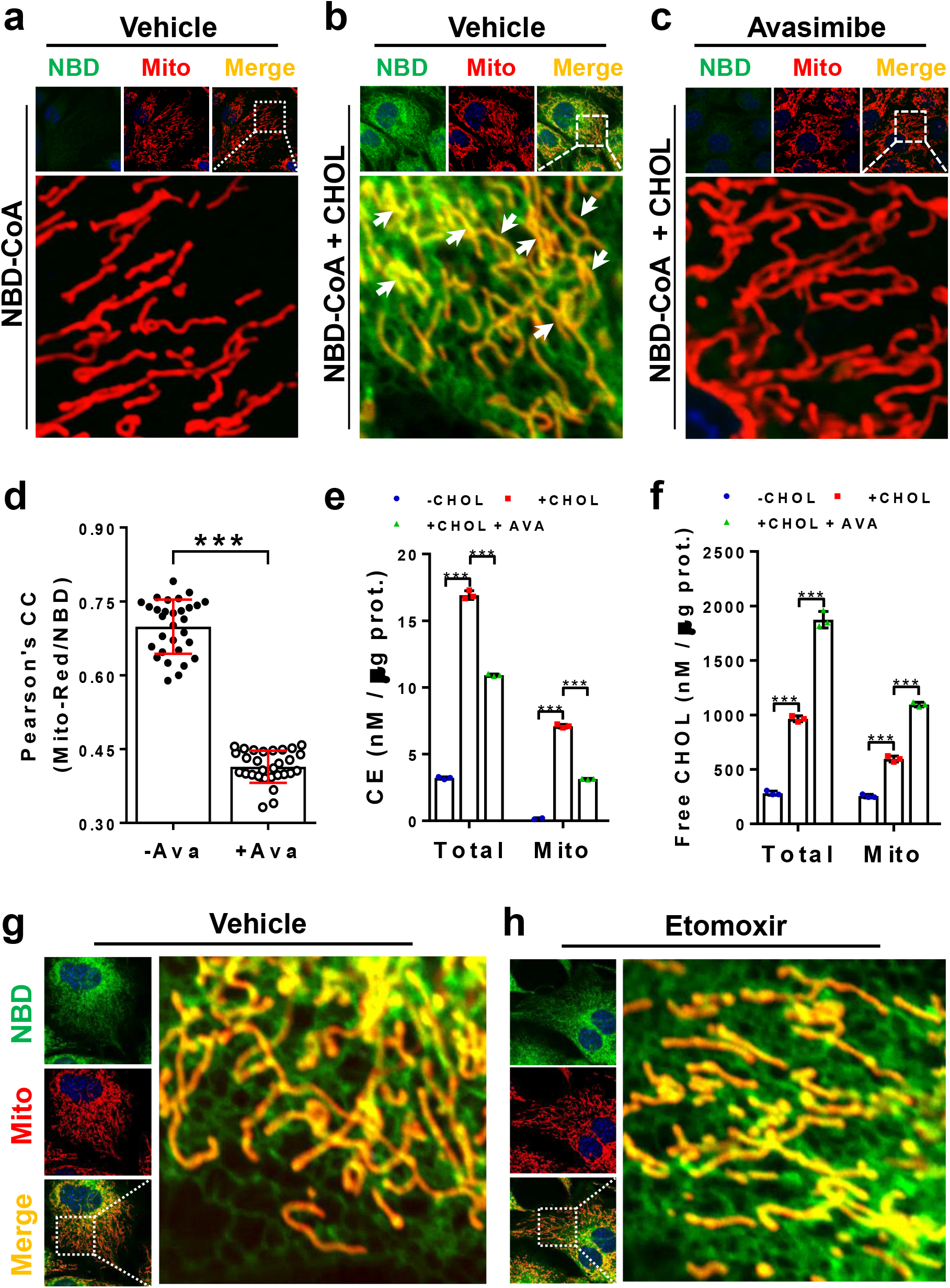
Labeling and confocal imaging of cholesterol esters in live cells. **a-b,** Confocal imaging analysis of the labeling and mitochondrial transport of cholesterol ester. C2C12 cells were starved in KRPH buffer for 1 hr, and then incubated with NBD-palmitoyl-CoA alone (**a**) or NBD-palmitoyl-CoA plus cholesterol (**b**) for 15 min. Mitochondria were stained with MitoTracker-Red. Arrows highlight the co-localization of NBD-lipid with mitochondria. **c,** Confocal imaging analysis of cholesterol esterification inhibition by treatment with avasimibe. C2C12 cells were pre-treated with avasimibe for 24 hr. Cells were then starved in KRPH buffer for 1 hr and then incubated with NBD-palmitoyl-CoA and cholesterol. **d,** Pearson’s correlation coefficient of mitochondria and NBD in the presence or absence of avasimibe. N=30 cells/group. **e**-**f,** Measurement of the total cell and mitochondrial cholesterol esters (CE, **e**) and free cholesterol (CHOL, **f**) in C2C12 cells. Cells pretreated with or without avasimibe were starved in KRPH buffer for 1 hr, and then incubated with palmitoyl-CoA alone (-CHOL) or palmitoyl-CoA plus cholesterol (+CHOL, or +CHOL + Avasimibe) for 30 min. N=3/group. **g-h,** Confocal imaging analysis of the labeling and mitochondrial transport of cholesterol ester in the presence (**h**) or absence (**g**) of etomoxir. C2C12 cells were pre-treated with or without etomoxir for 4 hrs, and then incubated with palmitoyl-CoA plus cholesterol for 15 min. Data are represented as mean ± SD, ***p<0.001 by Students t-test (**d**) or one-way ANOVA (**e-f**).

### Monoacylglycerol, but not diacylglycerol, promotes lipid droplet biogenesis

Triacylglycerol (TAG), which is composed of three fatty acyl groups esterified to a glycerol backbone at the *sn-1, sn-2* and *sn-3* positions, is used primarily for energy storage. Like phospholipids, TAG is also subjected to remodeling after *de novo* synthesis or after digestion in the gut, which is catalyzed sequentially by monoacylglycerol acyltransferases (MGAT) and diacylglycerol acyltransferases (DGAT)^17^. Using the lipid labeling method, we investigated the role of TAG remodeling in regulating lipid droplet biogenesis in live cells. COS-7 cells transiently expressing DsRed-ER5 were first stained with Bodipy 650/665, a red fluorescent dye for TAG, to label existing lipid droplets, then washed with culture medium, followed by treatment with NBD-palmitoyl-CoA alone or together with either monoacylglycerol (MAG) or diacylglycerol (DAG) as the substrates for the remodeling of TAG. As shown in Fig. 5, addition of MAG led to the labeling of nascent lipid droplets (LDs) which are still wrapped by ER sheets (Fig. 5a). The notion is supported by the observation that only a portion of MAG positive LDs were labeled by Bodipy 650/665 (Fig. 5a, arrows highlight the newly generated lipid droplets). In contrast, all DAG positive lipid droplets were labeled by Bodipy 650/665 (Fig. 5b, highlighted by arrows). The data suggest that the newly synthesized TAG from MAG was not only transferred to existing lipid droplets, but also stimulated the biogenesis of lipid droplets, whereas all the newly synthesized TAG from DAG was transferred to the existing lipid droplets in cells. Consistent with this notion, both MGAT1 and DGAT enzymes are localized at the ER, the primary site for lipid droplet biogenesis^18^, and overexpression of MGAT stimulates lipid droplet biogenesis^19^. In further support of the findings, the newly synthesized NBD-TAG can be detected by TLC analysis only in the presence of both NBD-palmitoyl −CoA and MAG or DAG (Extended Data Fig. 4d, highlighted by arrows).

**Fig. 5:**
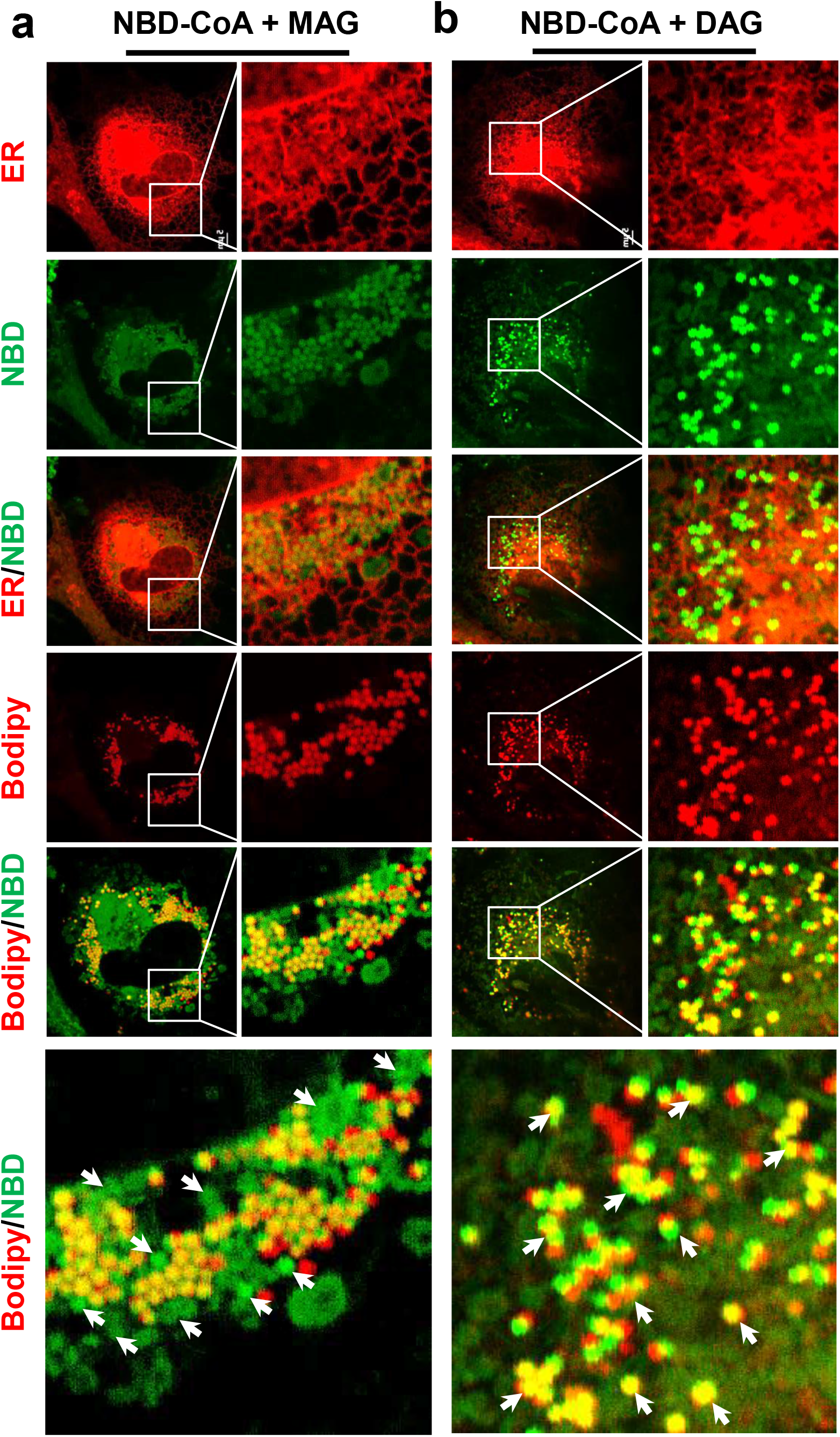
Labeling and confocal imaging of triacylglycerol in live cells. **a,** Confocal imaging analysis of COS-7 cells incubated with NBD-palmitoyl-CoA and 2-Oleoylglycerol (MAG). Arrows indicate the newly synthesized lipid droplets. **b,** Confocal imaging analysis of COS-7 cells incubated with NBD-palmitoyl-CoA and 1,2-Dioleoyl-sn-glycerol (DAG). Arrows indicate the co-localization of NBD-TAG with lipid droplets. COS-7 cells were starved in KRPH buffer containing Bodipy 650/665 to label lipid droplets for 1 hr before incubating with NBD-CoA plus MAG or DAG. ER was visualized by transfecting cells with DsRed-ER5.

### Newly remodeled phosphatidylethanolamine (PE) binds to autophagosomes during autophagy

To further confirm the newly NBD-labeled lipids were fully functional in cells, we next determined a role of PE remodeling in autophagosome biogenesis. During the initiation of autophagy, a cytosolic form of LC3 (LC3-I) is conjugated to PE to form LC3-PE conjugate (LC3-II), which is recruited to autophagosomal membranes^20^. We first tested the specificity of PE labeling in C2C12 cells incubated with NBD-palmitoyl-CoA and lysophosphatidylethanol (LPE). As shown in Fig. 6a, only a small portion of the *in vivo* labeled NBD-PE was co-localized with mitochondria, while most of them were localized in ER-like structures. Interestingly, much of the NBD-PE formed punctate structures, suggesting an association with microsomes such as autophagosomes (Fig. 6a, highlighted by arrows). To further investigate whether these NBD-PE punctas were co-localized with autophagosomes, COS-7 cells transiently expressing mRFP-LC3, a biomarker to visualize autophagosomes, were nutrient starved to initiate autophagy, and then incubated with NBD-palmitoyl-CoA and LPE. As shown in Fig. 6b, nearly all the newly remodeled NBD-PE punctas were co-localized with LC3 punctas, implicating a role of PE remodeling in autophagosome biogenesis. This notion is corroborated by a previous report that PE remodeling is stimulated by the onset of autophagy^21^.

**Fig. 6:**
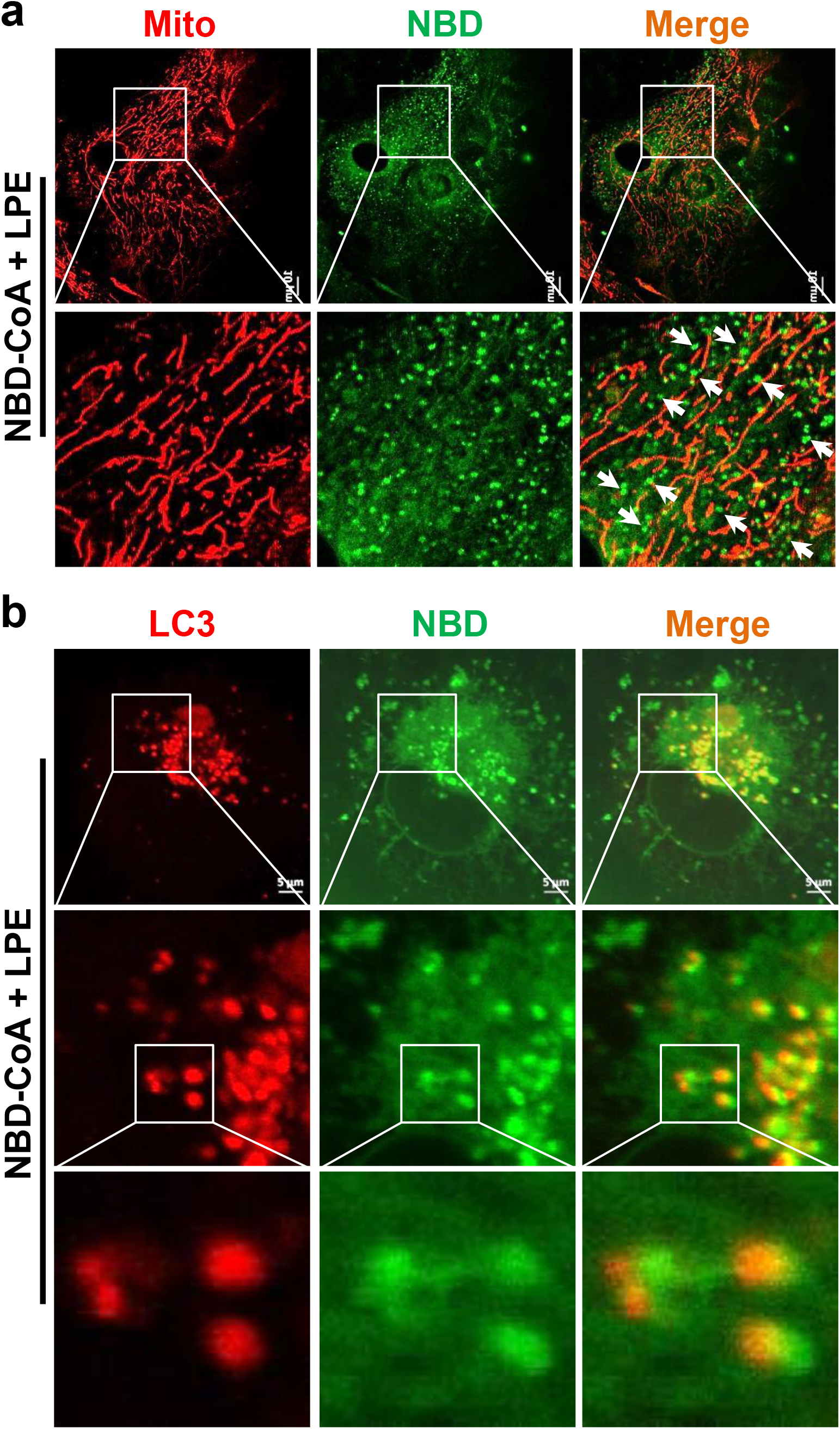
Labeling and confocal imaging of PE in live cells. **a,** Confocal imaging analysis of the labeling and mitochondrial localization of PE. C2C12 cells were starved in KRPH buffer for 1 hr and then incubated with NBD-palmitoyl-CoA and LPE for 15 min. Mitochondria were stained with MitoTracker-Red. Arrows indicate the NBD-PE punctas. **b,** Confocal imaging analysis of the co-localization of newly remodeled NBD-PE with autophagosomes. COS-7 cells transfected with mRFP-LC3 were starved in KRPH buffer for 1 hr, and then incubated with NBD-palmitoyl-CoA and LPE for 15 min.

### A striking difference between *de novo* remodeled lipids and exogenously labeled lipids in subcellular localizations

Exogenous fluorescence probe-labeled lipids are often used to investigate their trafficking in live cells by confocal imaging analysis^10, 22, 23^. However, it remains a major issue whether these exogenous fluorophore-labeled lipids are fully functional after loading into cells, since most of the exogenously labeled lipids end up in the plasma membrane and endosomes, possibly as a result of endocytosis and degradation^2^. We next investigated the subcellular localization of exogenous NBD-labeled phospholipids and cholesterol in live C2C12 cells stained with lysotracker Red, a fluorescent dye to label lysosomes. The results show that these exogenous NBD-labeled lipids, including NBD-PA, NBD-PS, NBD-PC, and NBD-cholesterol, demonstrate striking differences in subcellular localization with newly remodeled lipids. Unlike the *in vivo* NBD-labeled phospholipids through the remodeling pathway, almost all the exogenous NBD-labeled phospholipids and cholesterol co-localized with lysosomes (Fig. 7a-d), suggesting that these exogenous NBD-phospholipids and cholesterol are not fully functional in cells, and were primarily taken up by the lysosomes for degradation through lipophagy.

**Fig. 7:**
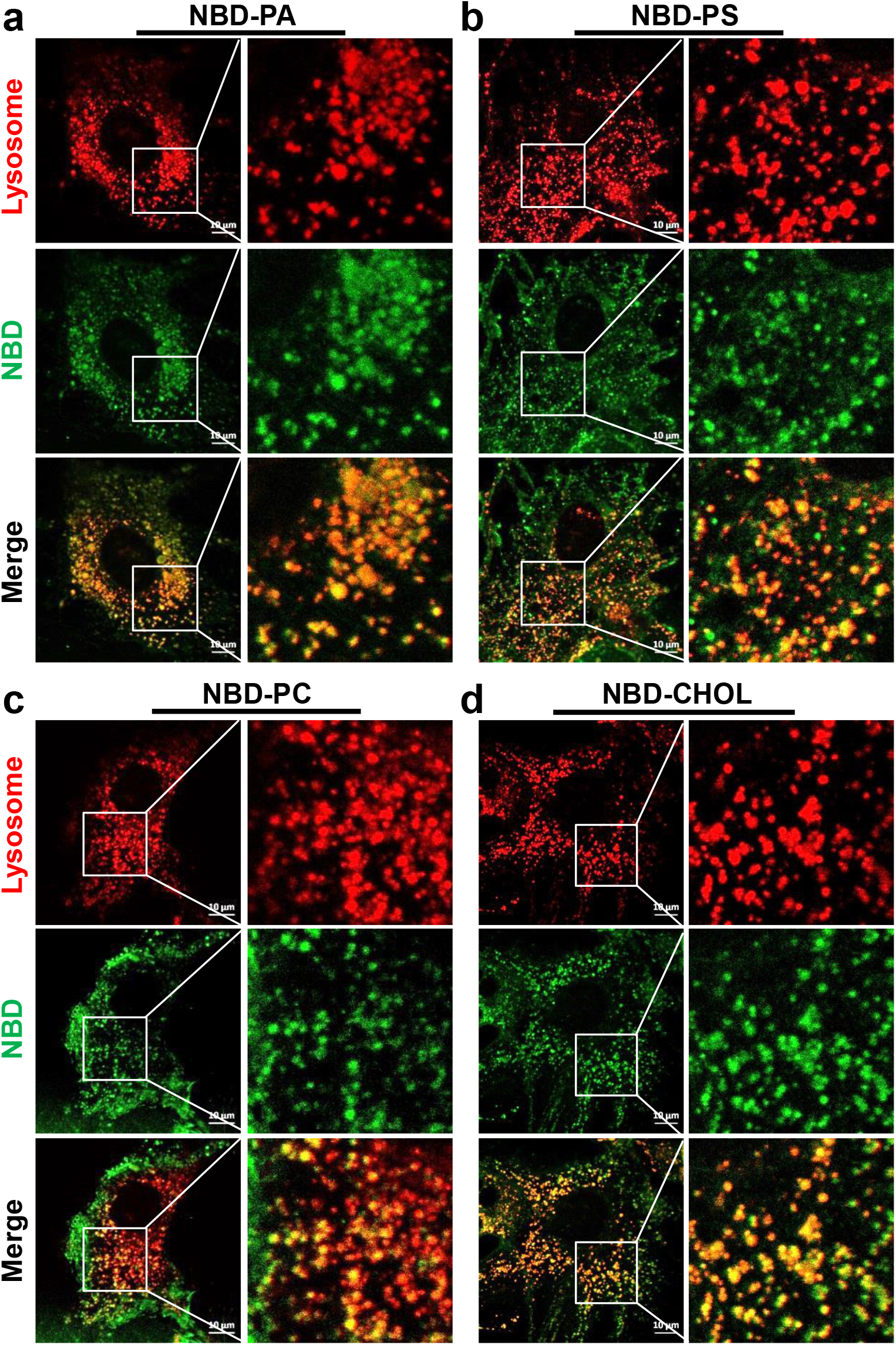
Subcellular localization of exogenous NBD-lipids in live cells. C2C12 cells were starved in KRPH buffer for 1 hr, and then incubated with exogenous NBD-lipids, including NBD-PA (**a**), NBD-PS (**b**), NBD-PC (**c**) or NBD-cholesterol (**d**). Lysosomes were stained with Lysotracker Red.

## Discussion

There are more than 1000 lipid species in a typical mammalian cell. Their level of complexity in subcellular organization and distribution within the cell is truly remarkable^24^. Therefore, it becomes a daunting task to investigate the spatiotemporal distribution of each individual lipid species. Microscopic visualization of lipids represents a powerful technique to study lipid behaviors in living cells. Since the vast majority of lipids are not intrinsically fluorescent, with the exception of a few lipid molecules such as dehydroergosterol and cholestatrienol^25^, various molecular probes were developed in recent years to visualize their subcellular localization, organization, and dynamics^2^. These probes include fluorophore-labeled lipids, antibodies, toxin domains, and genetically encoded protein domains^2^. However, none of the approaches can distinguish the acyl chain composition of any given lipid species *in vivo*, which is one of the most important features of a lipid. Additionally, there are many limitations with these approaches. For example, an attachment of a protein or large molecule to a lipid often disrupts its biophysical properties, leading to misplacement in subcellular distribution inside the cell^26, 27^. Moreover, these labeled lipids may not be fully functional, as evidenced by the observation that most of the exogenously labeled lipids end up in the lysosomes^5^, as confirmed by the current study which demonstrated a striking difference between *de novo* remodeled lipids and the exogenously labeled lipids in both subcellular localization and function. In contrast to *de novo* labeled lipids, the majority of the exogenously NBD-conjugated lipids, including NBD-PA, NBD-PC, NBD-PS, and NBD-cholesterol, end up in the lysosomes, possibly as a consequence of lipophagy.

Attachment of an NBD fluorescent group to the newly remodeled lipid molecules always raises a concern whether they faithfully represent the behavior of the endogenous lipids inside a living cell. We have addressed this important issue by multiple approaches. Firstly, we demonstrated that attachment of the NBD group to acyl-CoA did not affect its function as an acyl donor for the acyltransferase reaction, as evidenced by results from TLC analysis that the newly remodeled NDB-labeled lipids can only be detected if both NBD-acyl-CoA and lysophospholipids were supplemented in the culture medium. Secondly, we showed the attachment of the NDB molecule to the newly remodeled lipid molecules did not affect their recognition by the endogenous mitochondrial phospholipid transporters. Accordingly, ablation of the mitochondrial transporters prevented the mitochondrial trafficking of newly labeled NBD-phospholipids. Moreover, we demonstrated that newly remodeled NBD-tagged TAGs remain fully functional, as they were efficiently recruited by the lipid droplets. Furthermore, we demonstrated that the newly remodeled PE co-localized with autophagosomes, which is consistent with its projected role in the lipidation of the LC3 protein. Finally, we showed that inhibition of ACAT enzymes with a potent small molecule inhibitor completely abolished the production of NBD-labeled cholesterol esters and their transport to mitochondria.

Using this novel lipid labeling method, we not only validated the function of several mitochondrial phospholipid transporters and lipid metabolic enzymes, but also revealed some striking new functions of these transporters and enzymes, which was not possible by conventional biochemical assays. Consequently, we demonstrated that the TRIAP1/PRELI complex, which was previously reported to mediate mitochondrial PA transport^10^, also plays a key role in facilitating transport of PG from the ER to mitochondria. The findings are corroborated by a previous report that ablation of the TRIAP1 and PRELI protein complex caused apoptosis of cells, which can only be rescued by supplementation of excessive amount of PG^10^. The exogenous PG provided the substrate for PG remodeling through the Randel’s pathway. Additionally, we identified StARD7, a mitochondrial PC transporter^11, 28, 29^, as a novel regulator of mitochondrial PG transport. Our findings offered a logical explanation why depletion of StARD7 caused mitochondrial dysfunction^12^, since PG is a substrate for the synthesis of CL, and CL depletion caused both mitochondrial dysfunction and a loss of crista structure^30,31^. Furthermore, we also identified an unexpected role of ACAT enzymes in facilitating mitochondrial cholesterol transport. Accordingly, we showed that inhibition of ACAT enzymes by avasimibe, a potent inhibitor of both ACAT1 and ACAT2, not only prevented acylation of cholesterol, but also impaired mitochondrial transport of cholesterol esters. Since mitochondrial cholesterol transport is required for steroidogenesis, our findings offer an explanation why pharmacological inhibition of ACAT1 by a small molecule inhibitor significantly attenuated adrenocorticotropic hormone (ACTH)-stimulated cortisol production in an animal model of Cushing’s syndrome^32^. Finally, the lipid labeling method allowed us to identify an unexpected function of MAG in the biogenesis of lipid droplets. In contrast to DAG, treatment of cells with MAG not only led to the production of TAG, but also significantly stimulated lipid droplet biogenesis. The findings are consistent with our previous report that overexpression of the MGAT2 enzyme significantly stimulated lipid droplet biogenesis^19^.

Eukaryotic cells devote a substantial amount of energy to lipid synthesis, remodeling, and trafficking, which are mediated by a large number of enzymes and transporters. However, the functional importance of many of the enzymes and transporters remains poorly understood. Hence, we believe that our lipid labeling method can be used to speed up functional characterization of these enzymes and transporters. Additionally, lipid modification of proteins plays an important role in regulating protein trafficking and function. Acetylation and acylation regulate protein functions through diverse mechanisms, including stability, activity, subcellular localization, and crosstalk with other post-translational modifications^33^. However, it remains challenging to monitor the role of acetylation and acylation of a given protein in live cells. Although a photoactivatable probe was developed to investigate the lipids interaction proteins in cells, it has limitations in directly visualizing lipid localization and trafficking in living cell^34, 35^. Moreover, the *in vitro* conjugated photoactivatable lipids might be unlikely to mimic the function of the endogenous counterparts, as demonstrated by this study of the *in vitro* NBD labled lipids. Towards this end, our lipid labeling method can be easily adopted to investigate lipid protein interactions by using GFP or RFP-tagged proteins of interest. Although our current studies only tested NBD-acyl-CoA as the sole source of the green fluorescence, this principle can be easily adopted to other fluorescence-labeled acyl-CoA that emit red and blue colors, and thus will significantly expand the versatility of this lipid labeling method. Together with recent technological advances in microscopy, such as super resolution microscopy^36^, it can be envisaged this method will provide a powerful tool to investigate lipid remodeling, trafficking, and cellular functions in the context of diverse biological events inside a live cell.

## Acknowledgements

The current studies were funded in part by the NIH (DK076685, Y.S.), the American Diabetes Association (#1-18-IBS-329, Y.S.), the National Science Foundation of China (#31771309, Y.S.), and the National Institute on Aging (AG021890, J.N.)

## Author Contributions

Y.S. conceived the project and designed the research plan. J.Z., J.N., H.S. and J-P. A performed the experiments. Y.S., J.Z., J.N., H.S. and J-P. A. analyzed the data. Y.S. and J.Z. wrote the manuscript and all authors edited it.

## Methods

### Generation of gene knockout cells using CRISPR/Cas 9 gene editing

*Triap1, Prelid1 and StARD7* genes were knocked out in C2C12 cells by transfecting CRISPR/Cas9 mouse plasmids from Santa Cruz using Viafect Transfection Reagent (Promega E4982), and then purified through GFP and RFP fluorescence by the UT Health San Antonio Flow Cytometry Core. Cells were then kept under selection with 10 μg/mL puromycin (#sc-205821, Santa Cruz). Vector control cells were created by transfecting C2C12 cells with an empty pBABE-puro vector backbone (#1764, Addgene) and selected and kept in culture medium with 10 μg/mL puromycin.

### Labeling of newly remodeled phospholipids in cells

C2C12 or COS-7 cells were cultured in DMEM with 10% FBS and 1% penicillin/streptomycin. For lipid labeling, cells were first nutrient starved in a standard Krebs Ringer Phosphate HEPES buffer (KRPH, 140 mM NaCl, 2 mM Na_2_HPO_4_, 4 mM KCl, 1 mM MgCl_2_, 1.5 mM CaCl_2_, 10 mM HEPES, pH7.4) for 1 hr and then incubated with 16:0 NBD-CoA (1 μM, Avanti Polar Lipids, Cat# 810705) and lysophospholipids (20-100 μM, Avanti Polar Lipids) in KRPH buffer for 10-15 min. For visualizing of mitochondria, cells were either transfected with Mito-BFP, or stained with Mitotracker Red before nutrient starvation in complete medium (CM) for 15 min. For visualizing the endoplasmic reticulum (ER), cells were either transfected with DsRed2-ER5, or stained with ERtracker Red before nutrient starvation in CM for 15 min. Nuclei were stained by Hoechst 33342 (Invitrogene). Images were taken using a Zeiss LSM 710 confocal microscope.

### Labeling of newly remodeled cholesterol esters in cells

For labeling the newly remodeled cholesterol esters, C2C12 cells were first nutrient starved in KRPH buffer for 1 hr, and then incubated with 16:0 NBD-CoA (1 μM) and cholesterol (100 μM) in KRPH buffer for 10-15 min. For visualizing of mitochondria, cells were stained with Mitotracker Red before nutrient starvation in CM for 15 min. To inhibit the acylation of cholesterol in cells, C2C12 cells were first treated with avasimibe (10 μM), a selective acyl-CoA:cholesterol acyltransferase (ACAT) inhibitor, for 4 hrs in CM, and then subjected to mitochondrial staining and cholesterol labeling. Images were taken using a Zeiss LSM 710 confocal microscope.

### Labeling of newly synthesized triacylglycerol (TAG) in cells

COS-7 cells were transfected with DsRed2-ER5 to visualize the ER. After 36-48 hrs of transfection, cells were first nutrient starved in KRPH buffer for 1 hr, and then incubated with 16:0 NBD-CoA (1 μM) and 1,2-Dioleoyl-sn-glycerol (DAG, 25 μM) or 2-Oleoylglycerol (MAG, 25 μM) in KRPH for 10-15 min. To observe the co-localization of the newly synthesized TAG with lipid droplets, cells were first stained with Bodipy 650/665 in KRPH buffer for 1 hr during the nutrient starvation. Cells were washed twice with KRPH buffer, and then subjected to incubation with NBD-CoA and DAG or MAG. Images were taken using a Zeiss LSM 710 confocal microscope.

### Measurement of cholesterol and cholesterol ester contents

C2C12 cells were starved in KRPH buffer for 1 hr in the presence of vehicle or 10 μM avasimibe, and then incubated with 4 μM palmitoyl-CoA or 4 μM palmitoyl-CoA plus 50 μM cholesterol for 30 min. Cells were then harvested, and the mitochondria was fractionated and separated from the mitochondria associated membrane according to the method reported previously^37^. Total cell and mitochondrial free cholesterol and cholesterol ester contents were measured using the Cholesterol/Cholesterol Ester-Glo Assay Kit (J3190, Promega) according to the manufacturer’s instructions.

### Thin layer chromatography (TLC)

To confirm the remodeled or newly synthesized lipid products, cells were incubated with NBD-CoA and different lysophospholipids, DAG or MAG as indicated above. Total lipids were extracted using chloroform/methanol (2:1, v/v). The lipid samples were loaded to a TLC plate (Sigma), and developed in the solvent mixtures: chloroform: ethanol: water: trimethylamine (30:35:7:35, v/v/v/v) for phosphatidic acid (PA), or chloroform: methanol: water (65:25:4, v/v/v) for other phospholipids. For development of cholesterol esters, the lipid samples were loaded to the TLC plates which were first migrated in chloroform/methanol (1/1, v/v), and developed in the solvent mixture: hexane/diethyl ether/acetic acid (70/30/1, v/v/v). The lipid sample from cells only incubated with NBD-CoA was loaded as a negative control. The *in vitro* NBD labeled phospholipids were used as lipids markers. The TLC plates were dried after development and scanned using a Typhoon 9410 Scanner.

### Visualizing of *in vitro* NBD labeled lipids in cells

To observe subcellular localization of the *in vitro* NBD labeled lipids in cells, C2C12 cells were first nutrient starved in KRPH buffer for 1 hr, and then incubated with *in vitro* NBD labeled lipids (1 μM), including NBD-PA, NBD-PC, NBD-PS and NBD-Cholesterol, for 15 min in KRPH buffer. Lysosomes were stained with Lysotracker Red according to the manufacturer’s instruction. Images were taken using a Zeiss LSM 710 confocal microscope.

### Quantification and statistical analysis

Data were routinely represented as mean ± SD. Statistical significance was assessed by unpaired Student’s *t* test or one-way ANOVA using GraphPad Prism 6.0. Differences were considered statistically significant at p < 0.05; *p < 0.05; **p < 0.01; ***p < 0.001.

## Extended Data Figures and Legends

**Extended Data Fig. 1:**
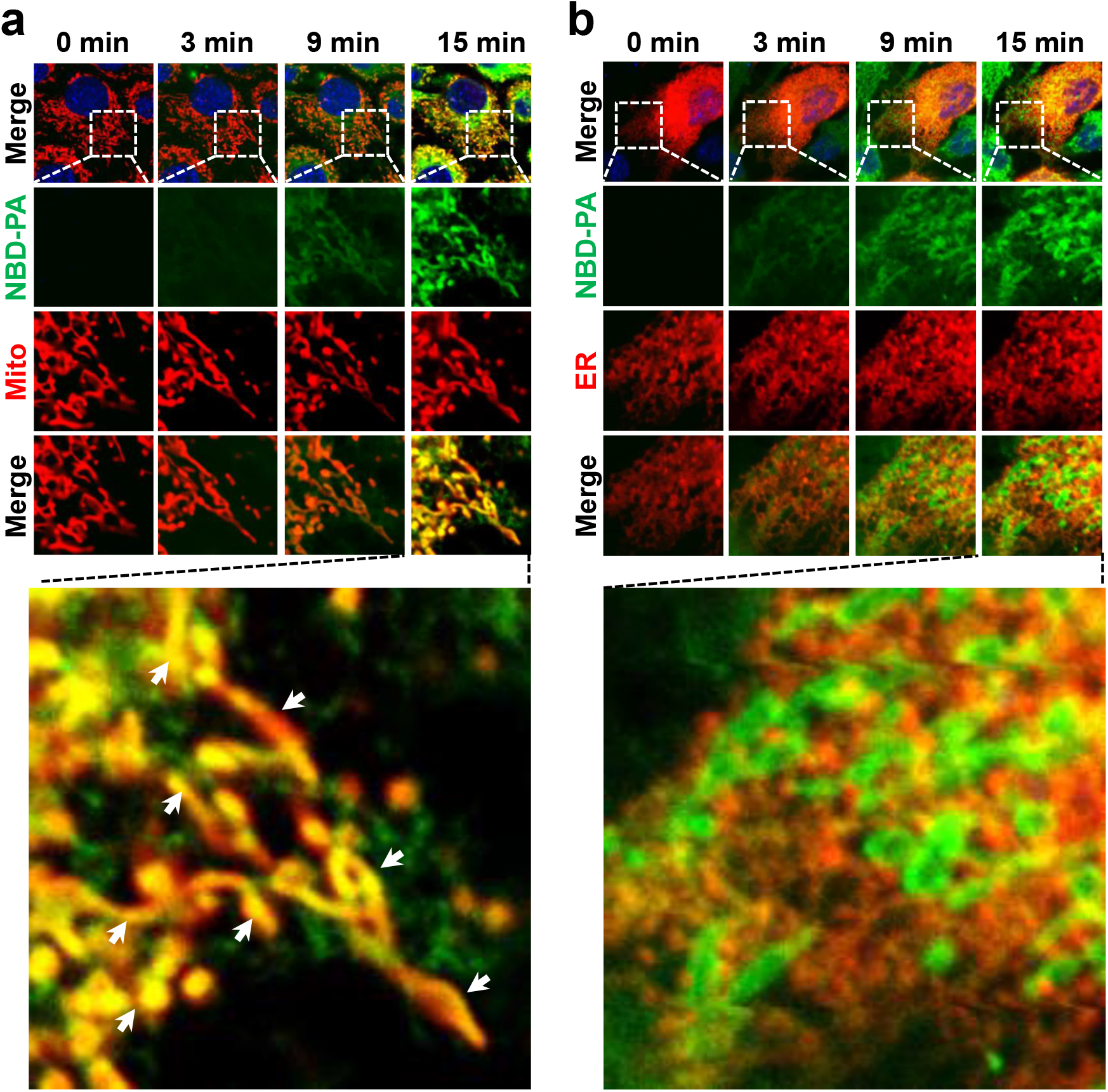
Time dependent labeling and mitochondrial transport of PA. **a,** Time-lapse confocal imaging analysis of NBD labeling and mitochondrial localization of PA in C2C12 cells. Arrowheads highlight the co-localization of NBD-PA with mitochondria. Arrows highlight the co-localization of NBD-PA with mitochondria. **b,** Time-lapse confocal imaging analysis of NBD labeling and the ER localization of PA in C2C12 cells. Cells were starved in KRPH buffer for 1 hr, and then incubated with NBD-palmitoyl-CoA and LPA. Images were taken at the indicated times after incubation. Mitochondria were stained with Mitotracker Red. ER was visualized by transfecting cells with DsRed-ER5.

**Extended Data Fig. 2:**
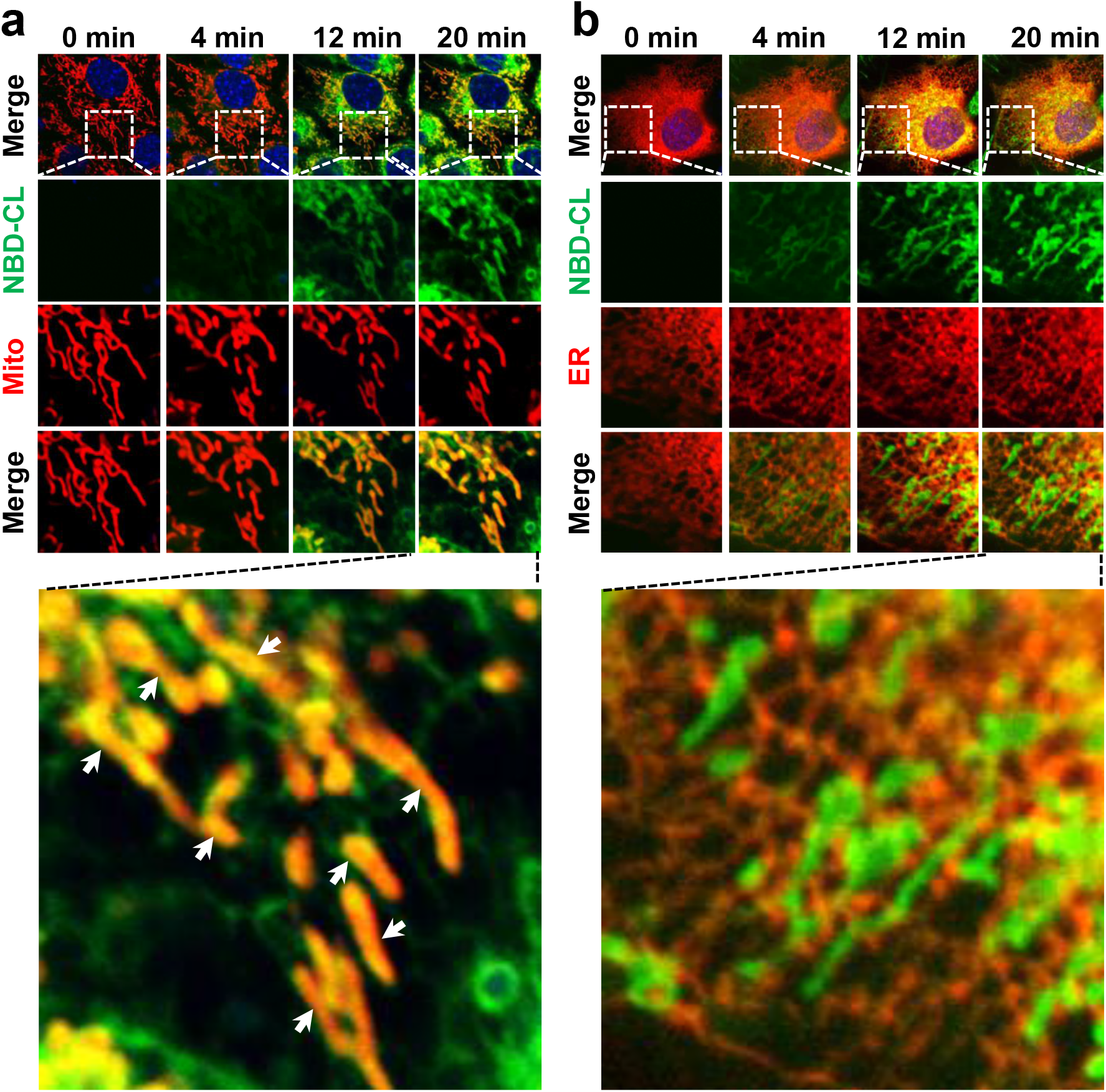
Time dependent labeling and mitochondrial transport of CL. **a,** Time-lapse confocal imaging analysis of NBD labeling and mitochondrial localization of CL in C2C12 cells. Arrow-heads highlight the co-localization of NBD-CL with mitochondria. Arrows highlight the co-localization of NBD-CL with mitochondria. **b,** Time-lapse confocal imaging analysis of NBD labeling and the ER localization of CL in C2C12 cells. Cells were starved in KRPH buffer for 1 hr, and then incubated with NBD-palmitoyl-CoA and MLCL. Images were taken at the indicated times after incubation. Mitochondria were stained with Mitotracker Red. ER was visualized by transfecting cells with DsRed-ER5.

**Extended Data Fig. 3:**
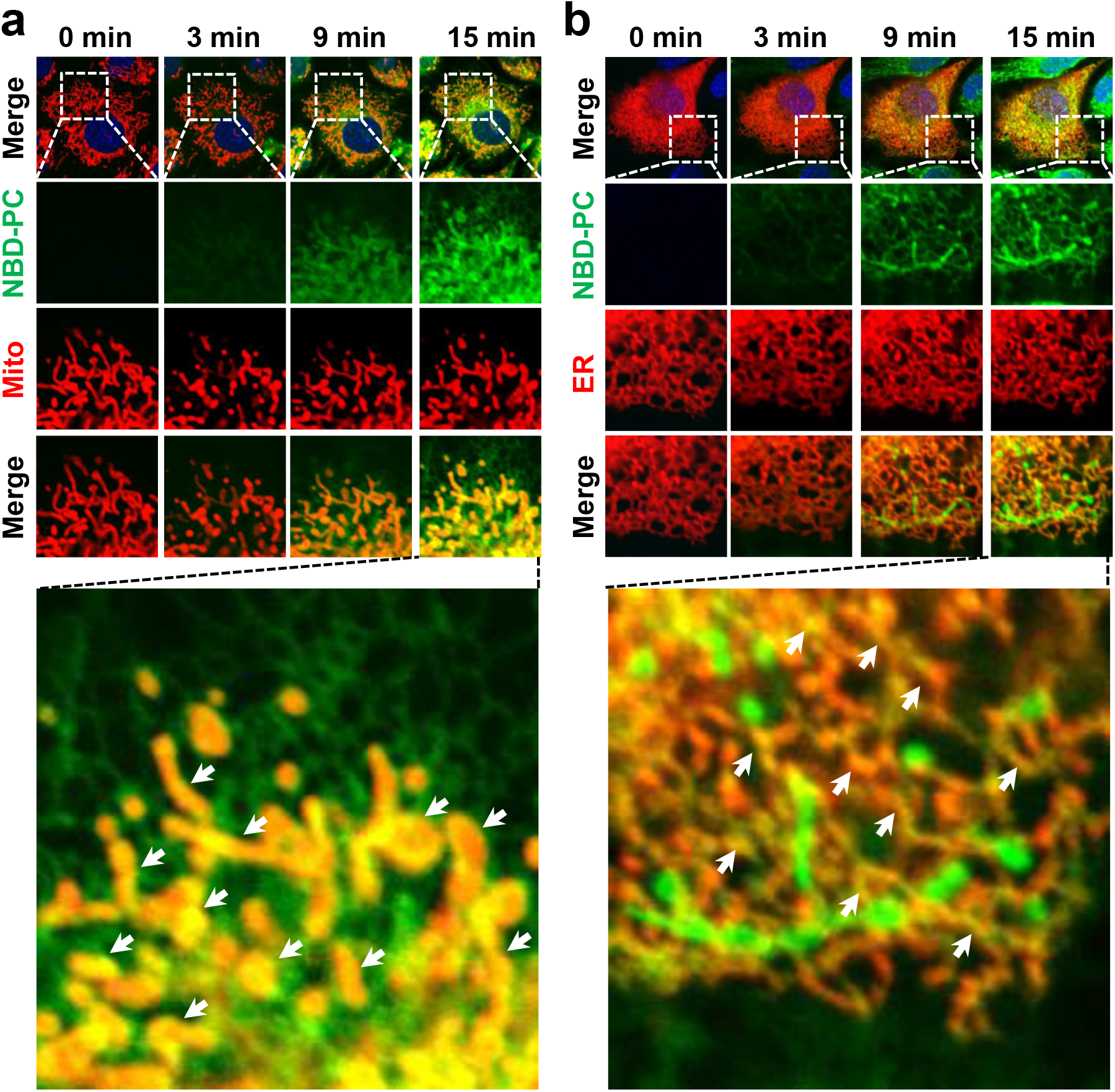
Time dependent labeling and mitochondrial transport of PC. **a,** Time-lapse confocal imaging analysis of NBD labeling and mitochondrial localization of PC in C2C12 cells. Arrows highlight the co-localization of NBD-PC with mitochondria. **b,** Time-lapse confocal imaging analysis of NBD labeling and the ER localization of PC in C2C12 cells. Arrows highlight the co-localization of NBD-PC with the ER. Cells were starved in KRPH buffer for 1 hr, and then incubated with NBD-palmitoyl-CoA and LPC. Images were taken at the indicated times after incubation. Mitochondria were stained with Mitotracker Red. ER was visualized by transfecting cells with DsRed-ER5.

**Extended Data Fig. 4:**
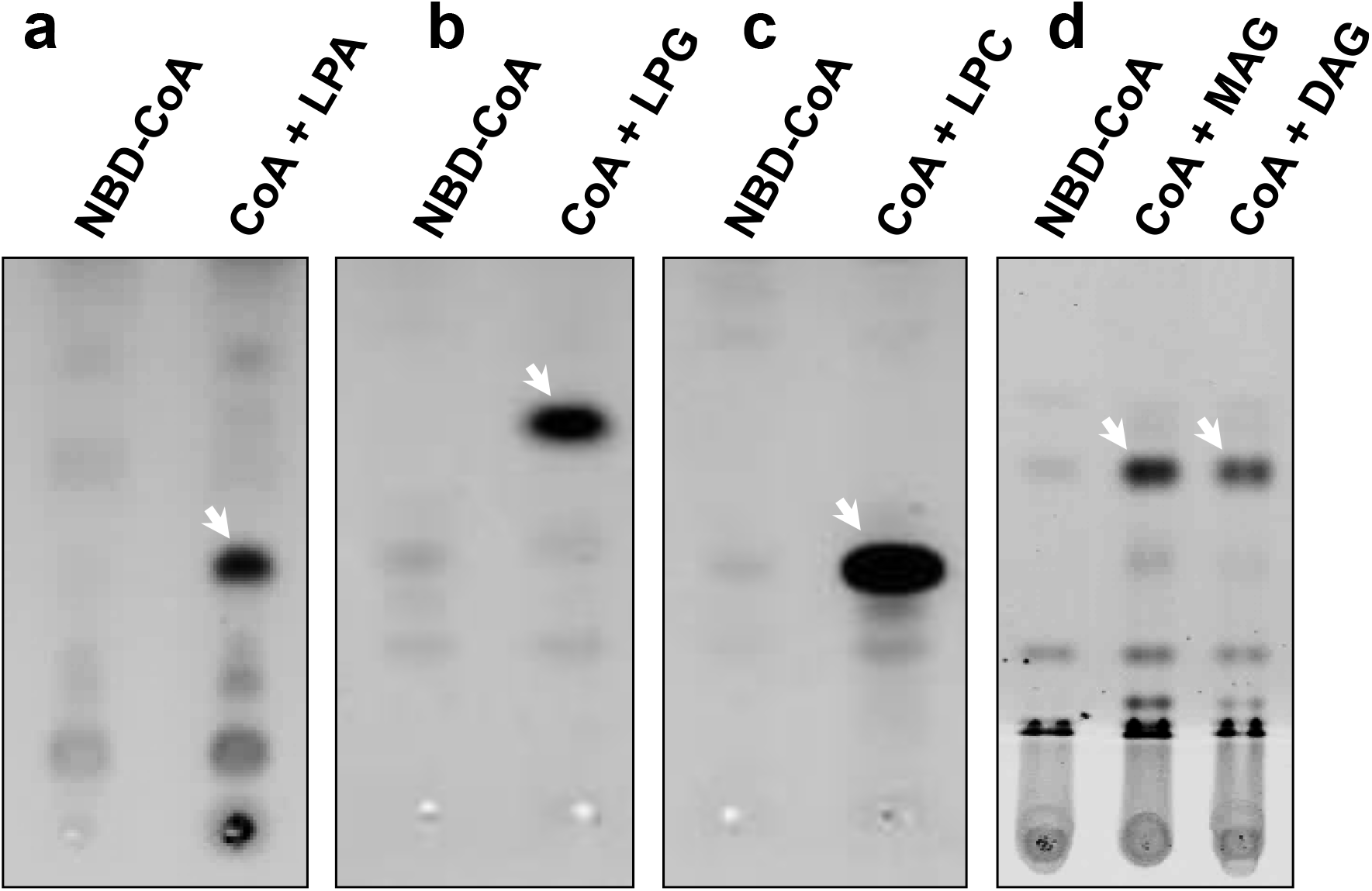
TLC analysis of NBD labeled lipids. **a-c,** TLC analysis of newly remodeled PA (**a**), PG (**b**) or PC (**c**) in C2C12 cells. Cells were starved in KRPH buffer for 1 hr, and then incubated with NBD-palmitoyl-CoA alone or with LPA, LPG or LPC for 15 min, respectively. **d,** TLC analysis of newly remodeled TAG in COS-7 cells. Cells were starved in KRPH buffer for 1 hr, and then incubated with NBD-palmitoyl-CoA and MAG or DAG for 15 min. The total lipids were extracted and developed by TLC as described in *Methods*.

**Extended Data Fig. 5:**
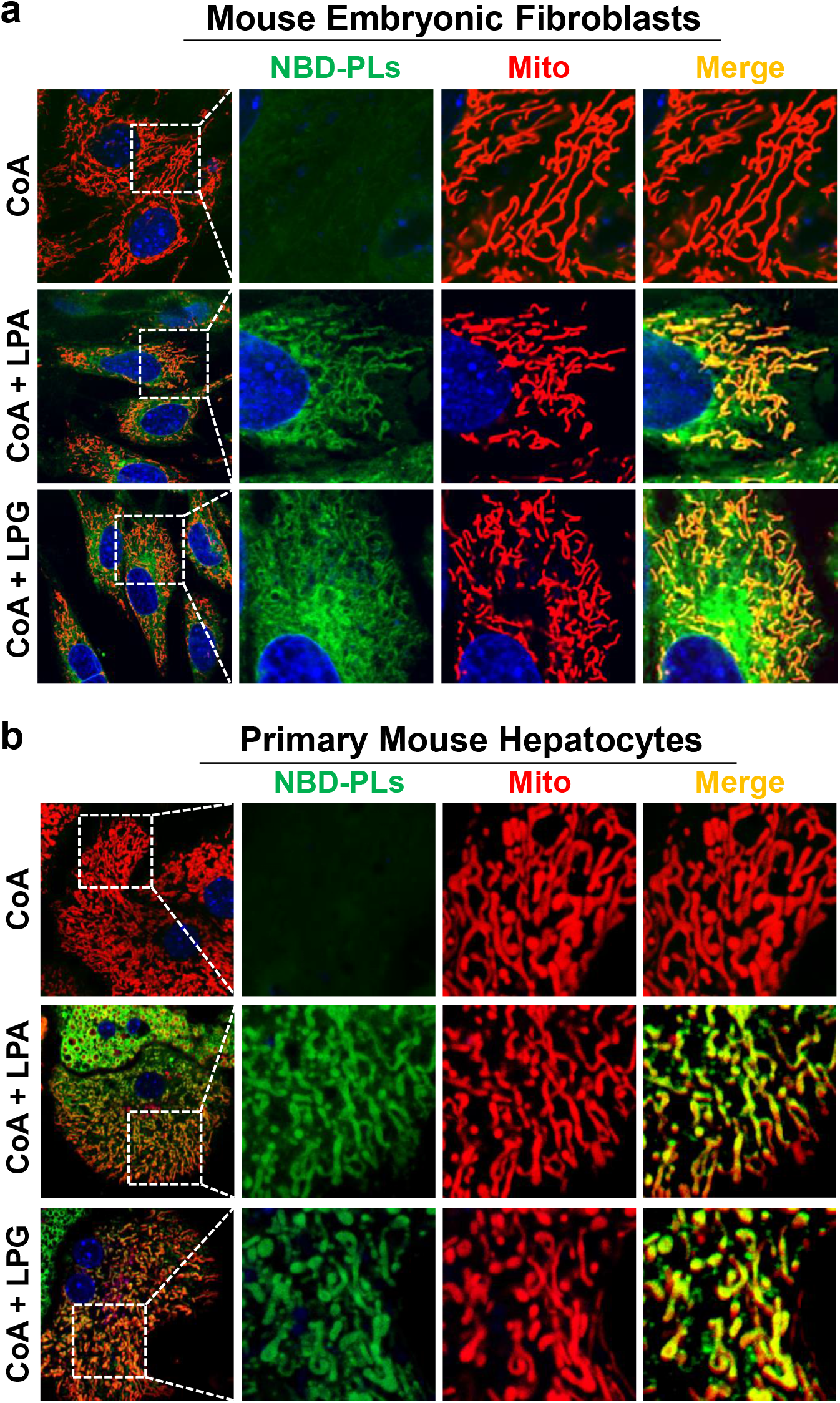
Labeling and confocal imaging of phospholipids in live primary cells. Primary mouse embryonic fibroblasts (**a**) and primary mouse hepatocytes (**b**) were starved in KRPH buffer for 1 hr, and then incubated with NBD-palmitoyl-CoA alone or NBD-palmitoyl-CoA plus LPA or LPG for 15 min. Mitochondria were stained with Mitotracker Red.

**Extended Data Fig. 6:**
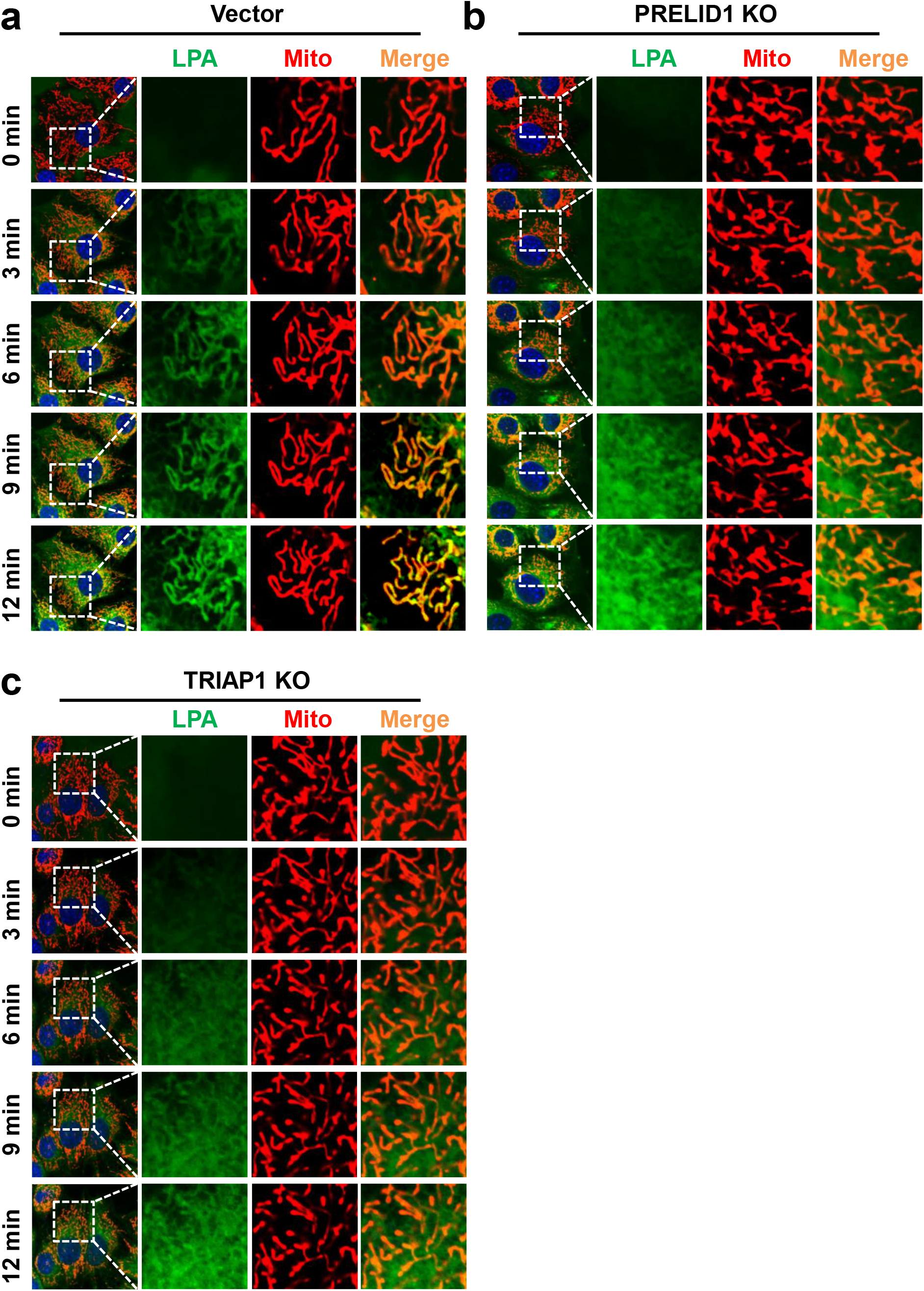
Time-lapse confocal imaging analysis of the labeling and mitochondrial transport of PA in live cells. **a-c,** Time-lapse confocal imaging analysis of the labeling and mitochondrial transport of PA in C2C12-vector control (**a**), PRELID1 KO (**b**) and TRIAP1 KO (**c**) cells. Cells were starved in KRPH buffer for 1 hr, and then incubated with NBD-palmitoyl-CoA and LPA. Images were taken at the indicated times after incubation. Mitochondria were stained with Mitotracker Red.

**Extended Data Fig. 7:**
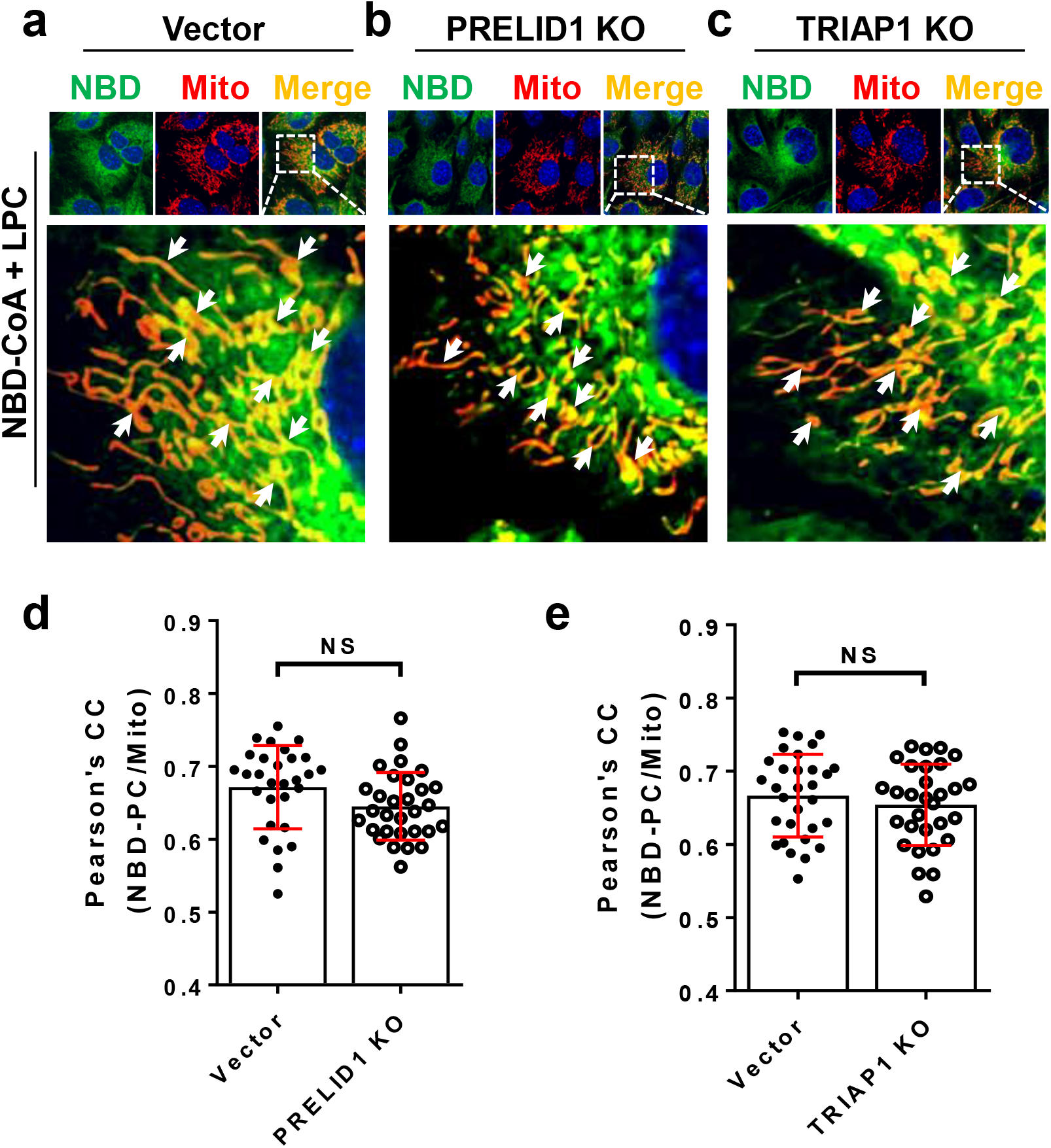
Labeling and confocal imaging of PC in PRELID1 and TRIAP1 KO cells. **a-c,** Confocal imaging analysis of the labeling and mitochondrial transport of PC in C2C12-vector control (**a**), PRELID1 KO (**b**) and TRIAP1 KO (**c**) cells. Arrows highlight the co-localization of NBD-PC with mitochondria. (**d-e**) Pearson’s correlation coefficient of mitochondria and NBD-PC in vector and PRELID1 KO (**d**) or TRIAP1 KO cells (**e**). n=30 cells/group. Cells were starved in KRPH buffer for 1 hr, and then incubated with NBD-palmitoyl-CoA and LPC for 15 min. Mitochondria were stained with Mitotracker Red. Data are represented as mean ± SD, ns, no significance by student’s t-test.

